# Clear Native Gel Electrophoresis for the Purification of Fluorescently Labeled Membrane Proteins in Native Nanodiscs

**DOI:** 10.1101/2025.03.21.644524

**Authors:** Bence Ezsias, Nikolaus Goessweiner-Mohr, Christine Siligan, Andreas Horner, Carolyn Vargas, Sandro Keller, Peter Pohl

## Abstract

Native gel electrophoresis techniques, such as blue or clear native gel electrophoresis (BNE or CNE), are widely used to separate and characterize proteins. However, in high-resolution CNE, mild anionic or neutral detergents are often used at concentrations too low to prevent membrane-protein aggregation. Additionally, the identification of proteins is hampered by the lack of suitable molecular-weight markers, like those used in SDS-PAGE. Here, we introduce a novel approach that combines charged polymer-encapsulated nanodiscs and fluorescence correlation spectroscopy (FCS) to address both challenges. Membrane proteins are first extracted using Glyco-DIBMA, a negatively charged amphiphilic copolymer. This enables the spontaneous formation of nanodiscs harboring the fluorescently labeled target protein within a native-like lipid-bilayer environment, which is confirmed by FCS. The nanodiscs are then subjected to detergent-free CNE. As the number of protomers increases, the nanodiscs grow larger resulting in increased migration distances in CNE due to higher charge densities. Crucially, the nanodiscs remain intact throughout CNE, as demonstrated by FCS analysis of resolubilized bands excised from the gels. Moreover, the membrane proteins used in this study: a potassium channel (KvAP), a sodium channel (NavMs), a water channel (GlpF), and a urea channel (*Hp*UreI) do not aggregate, as shown by the fluorescent brightnesses and diffusion times of individual nanodiscs. Moreover, the oligomeric states of membrane proteins can be deduced from the brightness per nanodisc. Since purified membrane proteins remain within a native-like lipid-bilayer environment and avoid detergent exposure, they are immediately suitable for downstream structural and functional studies.

## Introduction

Gel electrophoresis is an important analytical technique for separating proteins based on their molecular weight. Sodium dodecyl sulfate-polyacrylamide gel electrophoresis (SDS-PAGE) ^1^ remains one of the most widely used methods in cell and molecular biology. Since SDS provides a uniform charge-to-mass ratio for all protein species, their migration through the gel matrix depends solely on molecular weight. However, SDS’s strong denaturing properties disrupt not only protein–protein interactions but also the subunit connections within oligomeric structures. For electrophoretic analysis of intact, functionally active protein complexes, non-denaturing conditions are essential, while still shifting the intrinsic charge of the proteins. Electrophoresis on polyacrylamide gradient gels containing only 0.1% SDS was a first step in this direction ^2^. As the isolated photosynthetic complexes appeared green, this technique was named native green polyacrylamide gel electrophoresis and has been used extensively, often employing mild detergents like decyl maltoside (DM) or dodecyl glucoside (DDG) to solubilize cells and membranes of algae or higher plants ^3, 4^.

Another powerful non-denaturing method for gel electrophoresis is blue native PAGE (BNE). In this method, gradient polyacrylamide gels are used, but instead of mild detergents, the anionic triphenylmethane dye Coomassie Brilliant Blue G-250 (CBB) is employed to confer a net negative charge to the protein surface ^5^. BNE runs at a fixed pH of 7.5, allowing the separation of both membrane and soluble proteins regardless of their isoelectric point (pI) ^6^. Due to the charge shift, negatively charged proteins repel each other, reducing the likelihood of aggregation while maintaining the structure of protein complexes. However, BNE can be hindered by protein aggregation, particularly with membrane proteins ^7^.

In BNE and other gradient polyacrylamide gels, proteins are separated by molecular mass via a sieving effect ^8, 9^, with their migration toward the anode driven by their negative charge. The native structure of the protein influences its accessibility ^5, 6, 8-10^, making unambiguous identification challenging. Additionally, neutral detergents can compromise the oligomeric structure of sensitive protein complexes ^11^. Moreover, CBB can quench 90–95% of the fluorescence in labeled samples, which limits its utility in fluorescence-based assays ^12^.

A dye-free variant, clear native PAGE (CNE), addresses some of these challenges. Using the same buffer conditions as BNE but without CBB ^13^, CNE eliminates charge-shifting such that protein migration depends solely on the intrinsic charge of the protein. This lack of dye offers significant advantages for catalytic or fluorescent assays, but CNE is limited to acidic proteins (pI < 7); otherwise, proteins migrate toward the cathode and are lost ^14^. The absence of charged compounds also results in low resolution similar to BNE ^15^.

The use of small amounts of non-colored anionic detergents has improved CNE ^12^. High-resolution clear native PAGE (hrCNE) utilizes mixed micelles of sodium deoxycholate and dodecyl-β-D-maltopyranoside (DDM) in the cathode buffer. These buffer conditions impose a negative charge shift, maintain native protein conformation, and significantly increase resolution ^15^. However, even hrCNE struggles with membrane proteins due to persistent aggregation issues.

In native conditions like BNE and CNE, weight markers employed in SDS-PAGE cannot be used for accurate protein identification. Although commercially available sets of soluble protein markers exist ^6^, their use in native gel electrophoresis for membrane proteins is generally discouraged because conditions like running time and gel type can affect mass estimation ^6^.

A potential solution is the use of styrene/maleic acid (SMA) copolymers. They form nanodiscs that encapsulate membrane proteins with their native lipid bilayer, preventing misfolding and aggregation. ^16^. At the pH used in Tris-glycine PAGE (pH 8.3), deprotonation of the maleic acid residues gives SMA a high charge density, which far exceeds the inherent charge of the proteins and enables their migration ^17^. However, the SMA nanodiscs all have about the same size, irrespective of the size of the embedded membrane protein. Consequently, protein migration during electrophoresis is thought to be primarily determined by the size of the encapsulated proteins ^17^.

Here, we propose using a different type of charged polymer—Glyco-DIBMA ^18 19^, a partially amidated version of the diisobutylene/maleic acid copolymer DIBMA ^20, 21^. Glyco-DIBMA was introduced because its enhanced hydrophobicity makes it more efficient in nanodisc formation than other DIBMA copolymers ^18^. Additionally, Glyco-DIBMA offers an advantage over SMA that is crucial for the present purpose: the size of Glyco-DIBMA nanodiscs increases with the molecular weight of the encapsulated protein, allowing charge separation to differentiate between proteins of different sizes.

As an important additional improvement, we introduce the use of fluorescence to distinguish labeled membrane proteins and unambiguously identify their oligomeric states ^22^. Specifically, we use site-specific conjugation of fluorescent maleimide dyes to reduce cysteine residues via a thiol reaction after protein extraction into nanodiscs ^23, 24^. This allows visualization of mass-dependent nanodisc migration in the gel and further analysis of resolubilized protein bands using fluorescence correlation spectroscopy (FCS).^25^ FCS detects fluorescence intensity fluctuations in a diffraction-limited spot ^26, 27^, providing data on nanodisc residence time, molecular brightness (indicative of protein oligomeric state), and the concentration of labeled nanodiscs ^28^.

## Results

To demonstrate the utility of nanodiscs in the purification of membrane proteins, we selected four well-studied channels with known oligomeric states: a bacterial aquaporin (GlpF), which forms homotetramers, with each monomer containing a functional pore for water and glycerol conduction; an archaeal potassium ion channel (KvAP) and a bacterial sodium ion channel (*NavMs*), both of which form tetramers but contain only a single central functional pore; and a bacterial hexameric urea channel (*Hp*UreI) (*Fig. 1*).

**Figure 1.**
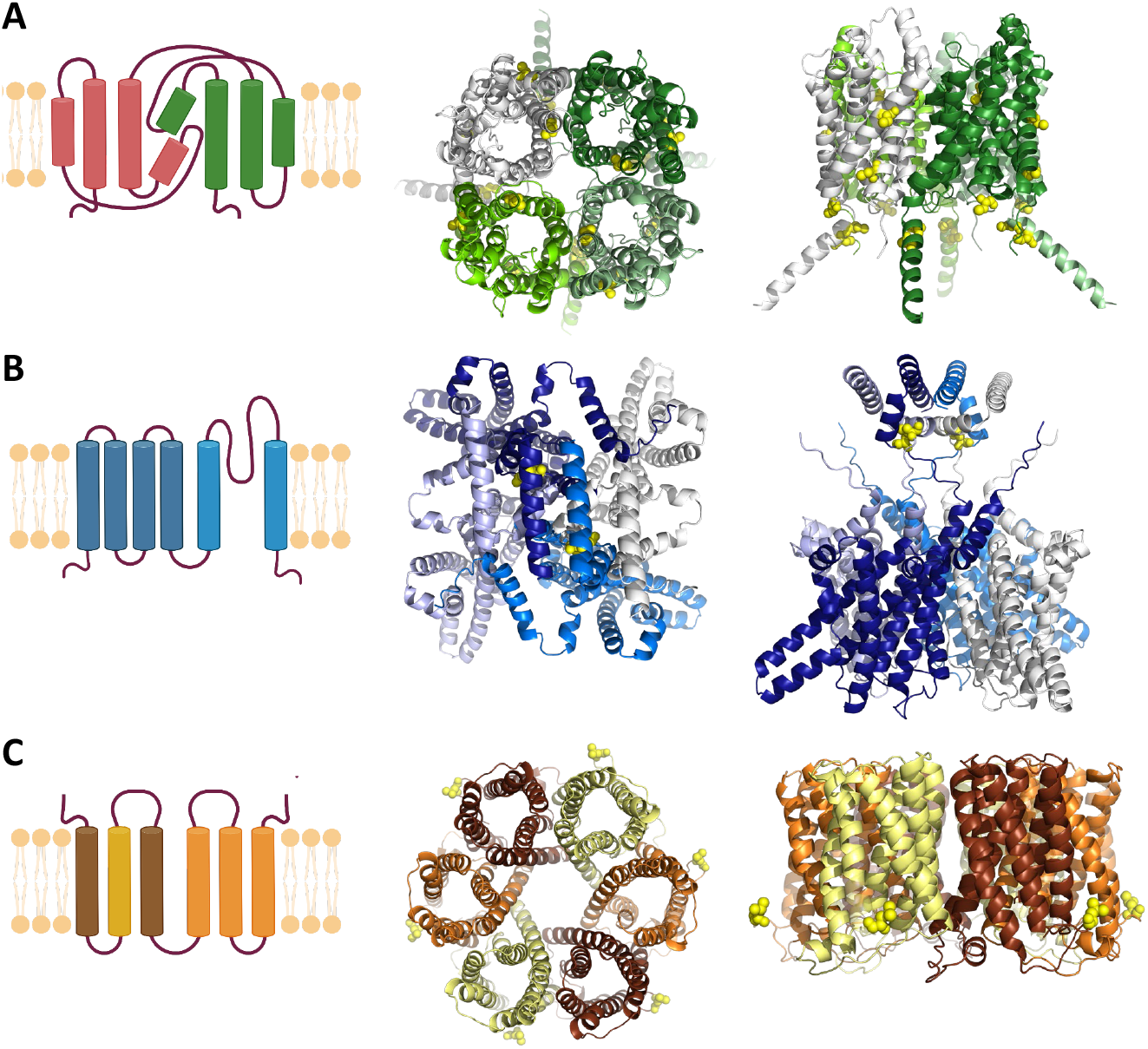
Schematic representation of membrane proteins with different folds, shown in both monomeric and oligomeric forms. The oligomeric models were generated using AlphaFold. This approach allows visualization of all labeling positions (yellow spheres), including those absent in the corresponding PDB structures. (A) Tetrameric glycerol uptake facilitator protein (GlpF) from Escherichia coli. All six cysteine positions are displayed, although not all can be labeled. (B) Tetrameric voltage-gated ion channels: KvAP from Aeropyrum pernix and NavMs from Magnetococcus marinus. Only the oligomeric model for KvAP is shown, with a single cysteine at position 260. (C) Hexameric urea channel (HpUreI) from Helicobacter pylori. L134C is the only available labeling position.

We started with the purification of the His-tagged homotetrameric glycerol uptake facilitator protein, GlpF. To this end, we added Glyco-DIBMA to the membrane fraction of *Escherichia coli* cells obtained after cell lysis by French press. The spontaneously formed nanodiscs were subjected to affinity chromatography (Fig. 2). We site-specifically labeled GlpF on-column with a maleimide dye, Alexa Fluor 647. The protein-containing elution fractions were then analyzed by fluorescence correlation spectroscopy (FCS) (Fig. S1), size exclusion chromatography (SEC), and blue native PAGE electrophoresis (BNE). GlpF-containing nanodiscs were extracted and resolubilized from the BNE bands and subjected to a second round of FCS analysis.

**Figure 2.**
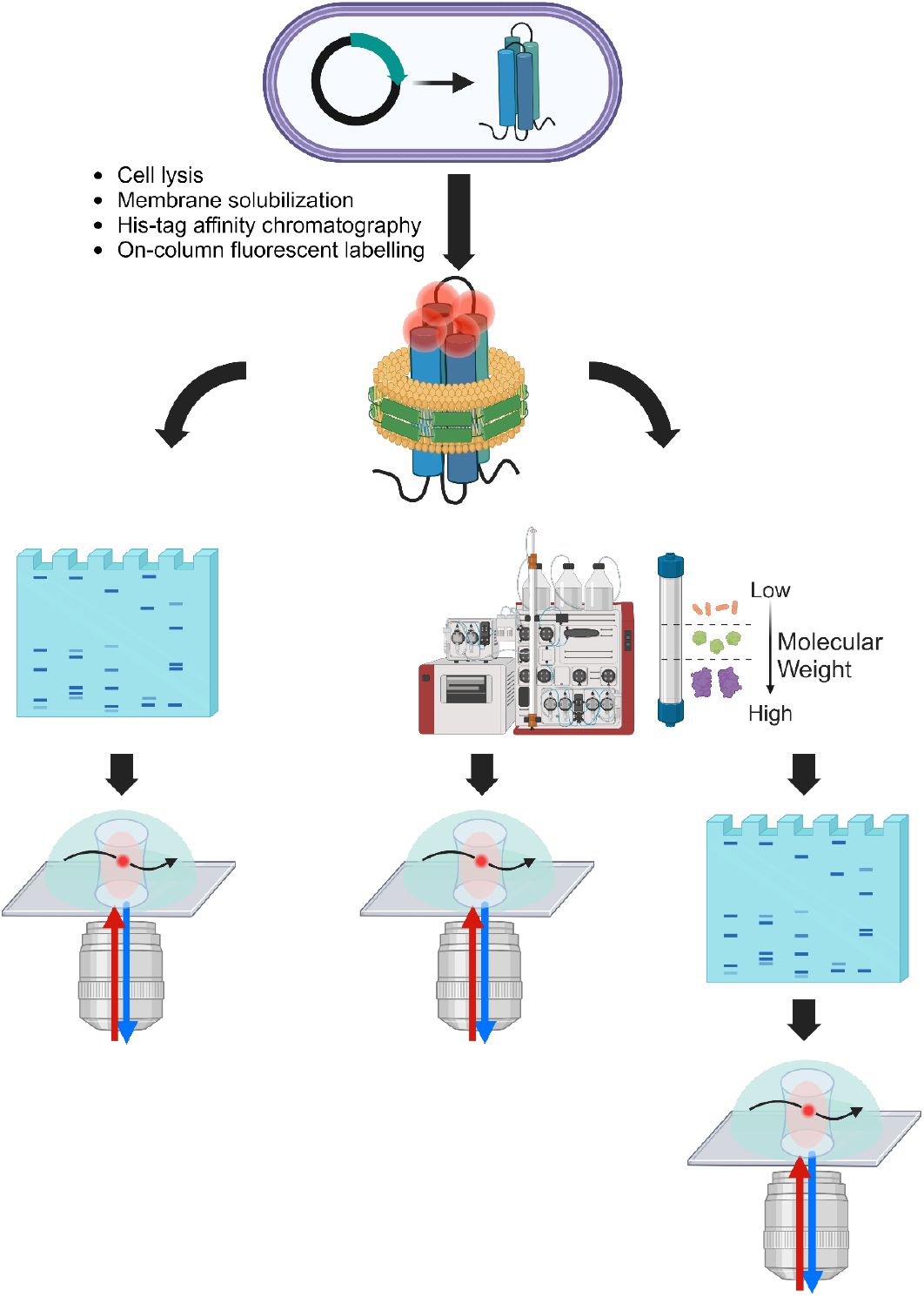
Schematic representation of the experimental workflow. Cell lysis and membrane solubilization by Glyco-DIBMA are followed by His-tag affinity chromatography and on-column labeling via thiol reaction. The eluted protein is then either directly subjected to native PAGE or first to size-exclusion chromatography and then to native PAGE. The protein samples extracted from the gels and purified only by size-exclusion are analyzed by FCS.

Both SEC and BNE revealed two major fluorescent fractions, along with several smaller, fainter ones (Figure 3). BNE allowed a rough size estimation of the nanodiscs but no unambiguous determination of the oligomeric state of the protein. The first peak in the SEC profile (Figure 3, B), appearing at an elution volume of 10–11 mL, is likely to represent the GlpF tetramer embedded in native nanodiscs. This tetrameric form should also correspond to the prominent fluorescent band on the BNE image (Figure 3, A). In contrast, the second peak in the chromatogram (Figure 3, A), at an elution volume of 15–16 mL, and the fainter fluorescent band on the BNE image (Figure 3, A) are likely to represent monomeric GlpF in nanodiscs.

**Figure 3.**
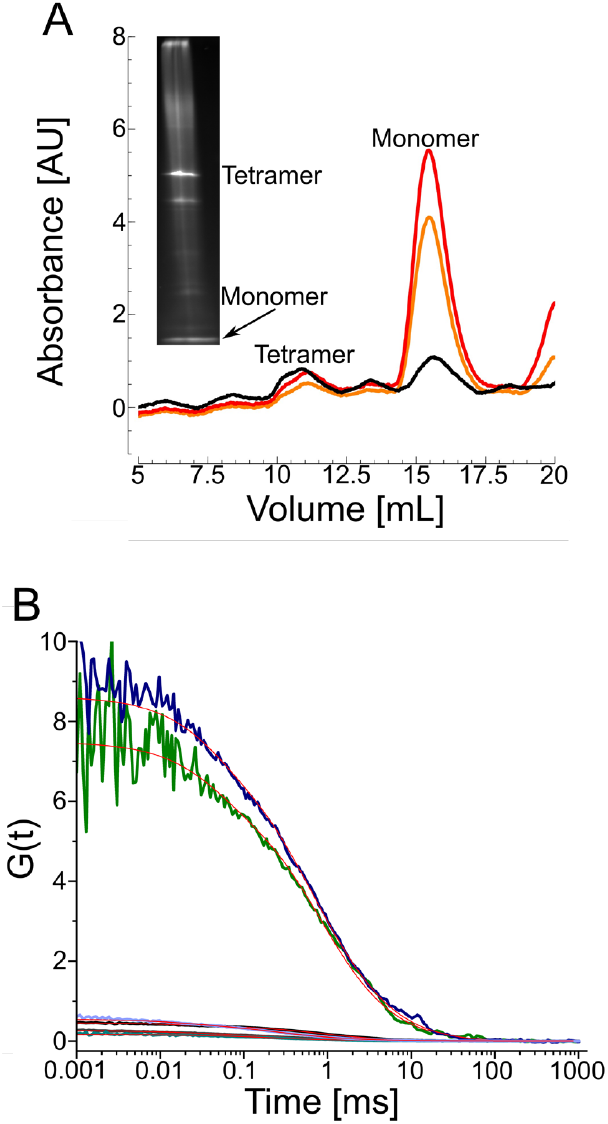
Blue native PAGE (BNE) compared to size-exclusion chromatography (SEC) and fluorescence correlation spectroscopy (FCS) of the glycerol uptake facilitator, GlpF. *A*: SEC and BNE fluorescent image, showing two major peaks, one for the tetramer (at 10–11 mL) and one for the monomer (at 15–16 mL). The black curve is the absorbance at 280 nm, the red at 650 nm, and the orange at 665 nm. *B*: Autocorrelation functions of tetrameric (black) and monomeric (grey) GlpF measured after SEC; after extraction from the total protein fraction and separation by BNE (blue for tetramer and purple for monomer); or after extraction from SEC-purified fractions and separation by BNE (green for tetramer and cyan for monomer). The estimated residence time and molecular brightness before BNE were 817±85 µs and 33±2 kHz for the tetramer and 360±72.6 µs and 8.5±0.15 kHz for the monomer. After BNE, these values were 854±240 µs and 7.6±0.9 kHz for the tetramer and 370±48.2 µs and 3.3±0.2 kHz for the monomer. The concentration of the extracted tetramer was 0.16 nM in 300 µL standard buffer.

Two rounds of FCS measurements—the first of SEC-purified samples and the second of resolubilized BNE bands support this interpretation (Figure 3, B). Specifically, we compared the residence times within the FCS focus and the molecular brightnesses per nanodisc (Table S1). For comparing molecular brightnesses, we used the brightness of 7.5 kHz measured for the free Alexa Fluor 647 maleimide dye as a ruler. We take the factor by which the particle brightness exceeds the ruler’s brightness in the first FCS round as the protein’s oligomeric state within the native nanodiscs. For GlpF, which is homotetrameric under native conditions, we measured molecular brightness of 8.5±0.15 kHz and 33±2 kHz before BNE. The former value corresponds to the monomer, while the latter represents the tetramer.

FCS also yields the hydrodynamic particle sizes (Table S2). Specifically, Eq. 1 (modified Einstein–Stokes equation) allows estimating the hydrodynamic particle diameter from the diffusion coefficient:

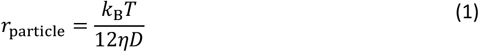

where *r*_particle_ is the hydrodynamic radius of the particle, *D* is the diffusion coefficient, *T* is the absolute temperature, *k*_B_ is the Boltzmann constant, and *η* is the viscosity.

We find hydrodynamic diameters of ∼15 nm for the native nanodiscs containing tetrameric GlpF and ∼6 nm for the native nanodiscs containing monomeric GlpF. Comparing these values to the diameter of the GlpF tetramer (7.9 nm) and monomer (4.0 nm), the purified nanodiscs appear sufficiently large to accommodate the different oligomeric forms of the protein. More interestingly, the GlpF tetramer appears to retain roughly 10 times more lipid molecules than the monomer (Table S3). Our observation of two oligomeric states of GlpF seems plausible since monomeric GlpF has been observed before in reconstituted vesicles ^29^.

The second round of FCS analysis, performed after BNE, yielded results consistent with those obtained before BNE: the diffusion time of the tetramer changed only slightly from 817±85 µs before BNE to 854±240 µs after BNE; while the diffusion time of the monomer changed from 360±72.6 µs before BNE to 370±48.2 µs after BNE. The consistent diffusion times indicate that the size of the nanodiscs was unaffected during BNE. However, the molecular brightness of the tetramer dropped from 33±2 kHz to 7.6±0.9 kHz, and that of the monomer from 8.5±0.15 kHz to 3.3±0.2 kHz. Obviously, the Coomassie dye dimmed the fluorescence of the Alexa dye. We also observed this effect for other purified membrane proteins, namely, the voltage-gated sodium ion channel NavMs, the potassium ion channel KvAP, and the urea channel *Hp*UreI.

### Nanodiscs and molecular brightness are preserved in clear native PAGE (CNE)

To mitigate the fluorescence quenching observed upon BNE, we substituted the Coomassie dye with mild detergents as commonly used in CNE, namely, 0.01% DDM and 0.05% sodium deoxycholate (DOC). Otherwise, the gel electrophoresis protocol remained unchanged. The purified and labeled prokaryotic voltage-gated sodium ion channel NavMs (Figure 4) and the potassium ion channel KvAP (Figure 5) were analyzed using high-resolution CNE after affinity chromatography and SEC. The SEC profiles of NavMs (Figure 4, A) exhibited two major fluorescent peaks corresponding to tetrameric and monomeric forms. CNE of the pooled protein elution displayed three fluorescent bands, with the highest band representing the NavMs tetramer (Figure 4, A).

**Figure 4.**
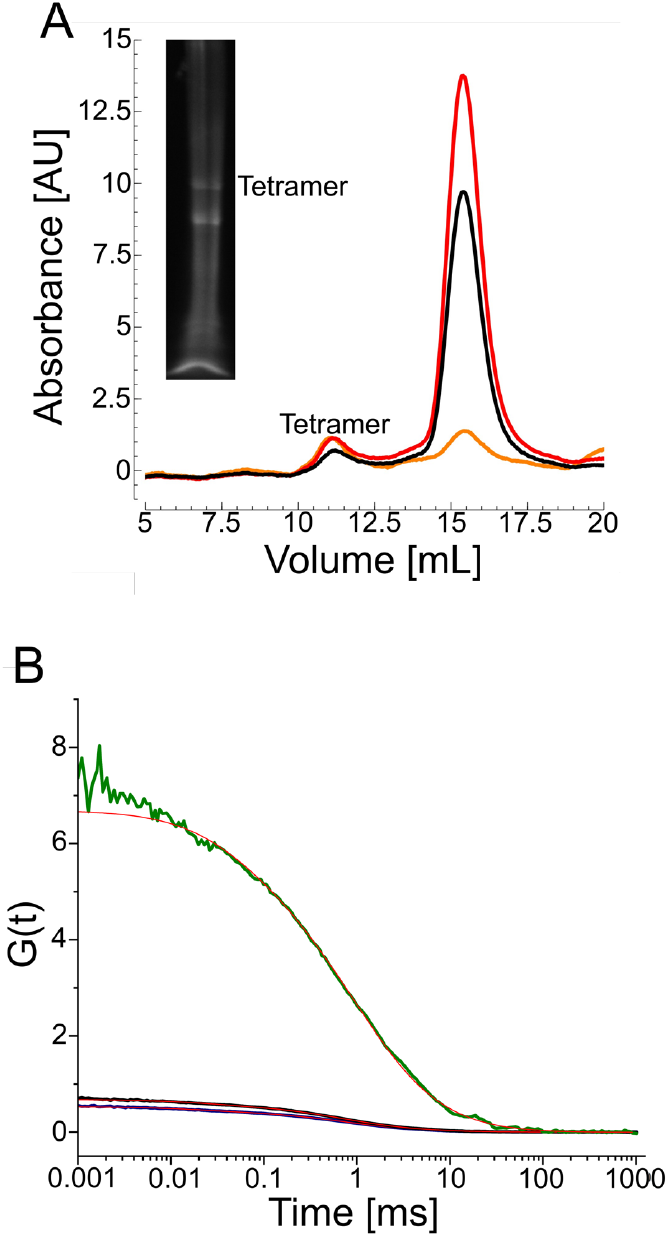
High-resolution clear native PAGE (CNE) compared to size-exclusion chromatography (SEC) and fluorescence correlation spectroscopy (FCS) of the voltage-gated sodium ion channel NavMs. *A*: SEC and high-resolution CNE fluorescent image of NavMs, labeled with Alexa Fluor 647 maleimide showing two major peaks and bands, one for a tetramer (at 10–11 mL) and one for smaller species (at 15–16 mL). The black curve is the absorbance at 280 nm, the red at 650 nm, and the orange at 665 nm. *B*: Fitted autocorrelation functions of tetrameric NavMs measured after SEC (black); after extraction from the total protein fraction and separation by high-resolution CNE (blue); or after extraction from SEC-purified fractions and separation by high-resolution clear CNE (green). The diffusion time and molecular brightness before CNE were 726±37 µs and 23.4±0.4 kHz, respectively. After CNE, these values were 740±62 µs and 21.5±0.9 kHz, respectively. The concentration of the extracted tetramer was 2.4 nM in 300 µL standard buffer.

**Figure 5.**
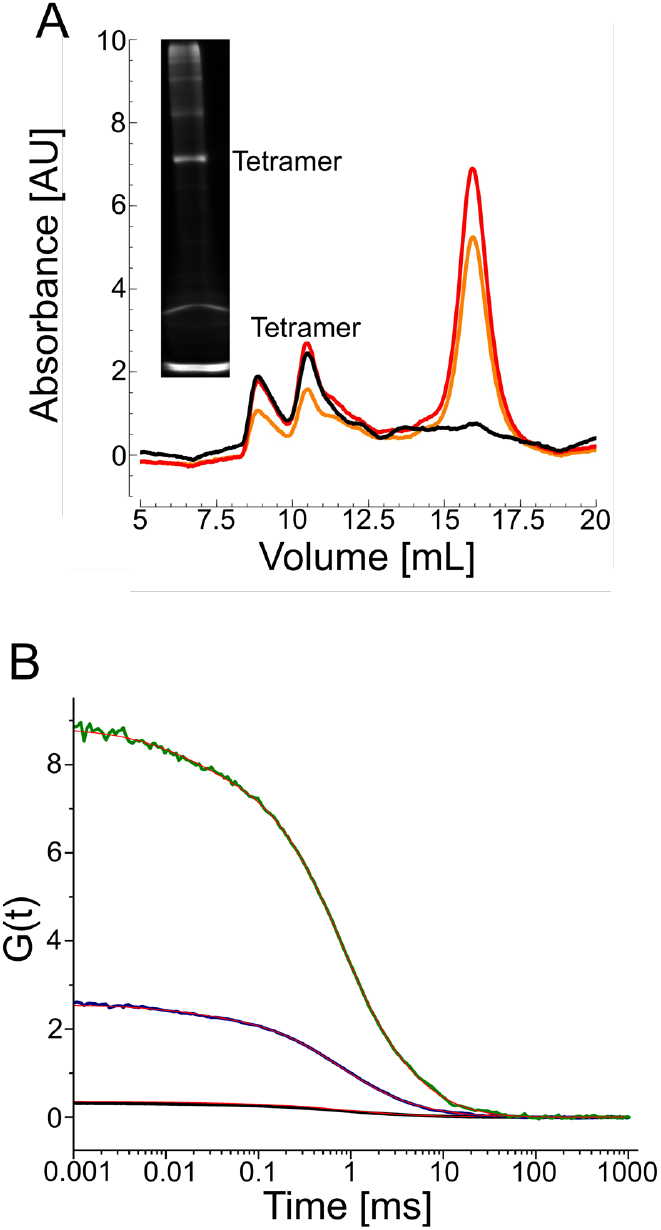
High-resolution clear native PAGE (CNE) compared to size-exclusion chromatography (SEC) and fluorescence correlation spectroscopy (FCS) of the voltage-gated potassium ion channel KvAP. *A*: SEC and high-resolution CNE fluorescent image of KvAP; SEC shows three major peaks, one indicating aggregate (at 8.5–9 mL), one for a tetramer (at 10–11 mL) and one for smaller species (at 15–16 mL). The black curve is the absorbance at 280 nm, the red at 650 nm, and the orange at 665 nm. *B*: Autocorrelation functions of tetrameric KvAP measured after SEC (black); after extraction from the total protein fraction and separation by high-resolution CNE (blue); or after extraction from SEC-purified fractions and separation by high-resolution CNE (green). The diffusion time and molecular brightness before CNE were 880±64.1 µs and 44±0.5 kHz, respectively. After CNE, these values were 770±107 µs and 48±1.7 kHz, respectively. The concentration of the extracted tetramer was 0.43 nM in 300 µL standard buffer.

FCS measurements of NavMs (Figure 4, B) conducted before CNE indicated an average count of 23.4±0.4 kHz for the SEC-purified tetramer. The corresponding residence time in the confocal volume was 726±37 µs. This yielded a hydrodynamic diameter of 17.52 nm for the protein-containing nanodiscs, which readily accommodates a tetrameric protein with a diameter of 6 nm. FCS measurements after CNE revealed an essentially unchanged molecular brightness and residence time of 21.5±0.9 kHz and 740±62 µs, respectively. The ratio of molecular brightness between the NavMs tetramer and the ruler is 3-fold; the deviation from the expected value of 4 could be due to self-quenching between the fluorescent labels on the monomers constituting a tetramer.

In both SEC and CNE, a second NavMs fraction appeared as well. Although it is unclear if this fraction represents a smaller oligomer—such as a monomer or a dimer—embedded within nanodiscs or rather some other kind of nanoparticles, the ratio of molecular brightness between this fraction and the ruler is ∼1.8-fold, indicating a potential dimeric form of NavMs. Both dimer (4.5 nm in diameter) and monomer (2.2 nm in diameter) would readily fit into the particles in question, which have a hydrodynamic diameter of ∼6 nm.

We made similar observations with the purified KvAP channel (Figure 5). Both SEC and CNE revealed two major fluorescent fractions corresponding to the tetrameric and monomeric forms of the protein (Figure 5, A). FCS measurements (Figure 5, B) yielded residence times of 880±64.1 µs before CNE and 770±107 µs after CNE. The molecular brightness values were 44±0.5 kHz and 48±1.7 kHz, respectively. The molecular brightness of KvAP was 5.5-fold higher than that of the ruler and, thus, significantly greater than the value of 4 expected for a tetrameric protein. This can be explained by the observation that surface hydrophobicity may affect the quantum yield ^30^. Here, the diameter of the native nanodiscs amounted to ∼23 nm, while the KvAP tetramer has a diameter of 10.5 nm. As for NavMs, SEC and CNE both displayed a second fluorescent peak, with a residence time similar to that of the ruler. As for GlpF, this finding indicates a monomer embedded in nanodiscs. We determined the hydrodynamic size of this fluorescent particle to be ∼7 nm, while the KvAP monomer has a diameter of 4.5 nm.

### CNE of protein-containing nanodiscs does not require the presence of detergent

Since Glyco-DIBMA is electrically charged, we wondered whether the addition of a charged detergent could be avoided. To test this hypothesis, we subjected the same KvAP-containing elution fraction obtained from SEC to detergent-free CNE, yielding results similar to those obtained with detergent (Figure 6). Specifically, KvAP migrated identically in both gels, with two bright bands indicating predominantly tetrameric protein and a smaller monomeric fraction (Figure 6, A). This observation demonstrates that Glyco-DIBMA conferred a charge density to the native nanodiscs that is sufficient to enable their migration towards the anode. SEC further validated the consistency between high-resolution CNE and simple, detergent-free CNE, revealing a tetramer peak (Figure 6, A) reminiscent of that observed in high-resolution CNE (Figure 5, A). FCS confirmed the tetrameric state of nanodisc-embedded KvAP, showing a 4-fold increase in molecular brightness compared to the ruler and the same particle diameter as measured after detergent-based CNE. Moreover, the diffusion times of the tetrameric fractions obtained from high-resolution, detergent-based CNE and simple, detergent-free CNE were similar, both ranging between 700 µs and 800 µs (Figure 6, B).

**Figure 6.**
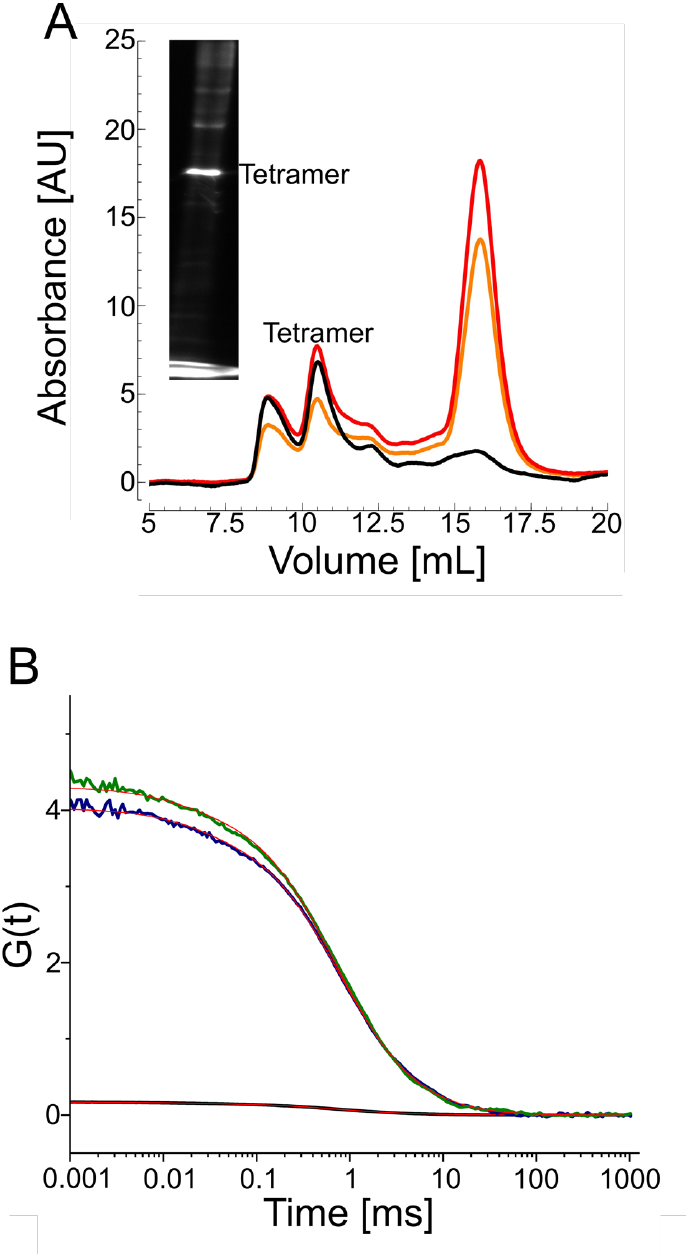
Detergent-free clear native PAGE (CNE) compared to size-exclusion chromatography (SEC) and fluorescence correlation spectroscopy (FCS) of the voltage-gated potassium ion channel KvAP. *A*: SEC and detergent-free CNE fluorescent image of KvAP; SEC shows three major peaks, indicating aggregates (at 9 mL), tetramers (at 10–11 mL), and a smaller species (at 15–16 mL). The black curve is the absorbance at 280 nm, red at 650 nm, and the orange at 665 nm. *B*: Autocorrelation functions of tetrameric KvAP measured after SEC (black); after extraction from the total protein fraction and separation by detergent-free CNE (blue); or after extraction from SEC-purified fractions and separation by detergent-free CNE (green). The diffusion time and molecular brightness before CNE were 747±52.6 µs and 43±1.8 kHz, respectively. After CNE, these values were 760±20 µs and 38±1.5 kHz, respectively. The concentration of the extracted tetramer was 0.27 nM in 300 µL standard buffer.

Finally, to demonstrate the versatility of detergent-free CNE, we applied it to a membrane protein of a completely different fold, namely, the homohexameric urea channel *Hp*UreI (Figure 7). The hexameric and monomeric forms of the protein embedded in native nanodiscs were separated by SEC (Figure 7, A), and their diameters were determined by FCS to be 16.86 nm and 6.7 nm, respectively. For comparison, the *Hp*UreI hexamer and monomer have protein diameters of 9.3 nm and 4.1 nm, respectively. FCS measurements before and after detergent-free CNE revealed no changes in residence time or molecular brightness. The residence times were 811±58.7 µs and 830±84.2 µs before and after CNE, respectively (Figure 5, B), while the molecular brightness was 36±1.7 kHz before CNE and 37±0.6 kHz after CNE, both in excellent agreement with the 6-fold increase over the brightness of the ruler.

**Figure 7.**
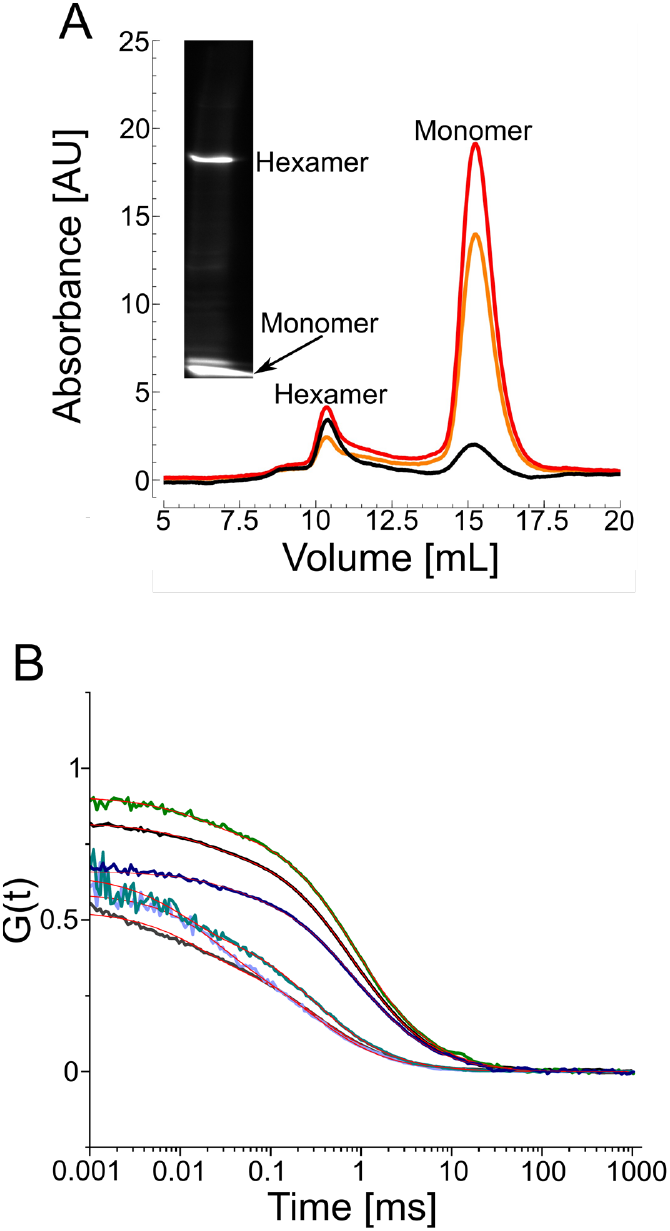
Detergent-free CNE of the urea channel HpUreI embedded in native nanodiscs compared to size-exclusion chromatography (SEC) and fluorescent correlation spectroscopy (FCS). *A*: SEC and CNE fluorescent image of the total protein elution *Hp*UreI; SEC shows two major peaks, one for a hexamer (at 10–11 mL) and one for a monomer (at 15–16 mL). The black curve is the absorbance at 280 nm, the red at 650 nm, and the orange at 665 nm. *B:* Autocorrelation functions of hexameric (black) and monomeric (grey) *Hp*UreI, measured after SEC; after extraction from the total protein fraction and separation by CNE (blue for the hexamer and purple for the monomer); or after extraction from SEC-purified fractions and separation by CNE (green for the hexamer and cyan for the monomer). The diffusion time and molecular brightness before CNE were, respectively, 811±58.7 µs and 36±1.7 kHz for the tetramer and 320±32.5 µs and 9±0.5 kHz for the monomer. After CNE, these values were, respectively, 830±84.2 µs and 37±0.6 kHz for the tetramer and 270±9.6 µs and 7±0.2. kHz for the monomer. The concentration of the extracted hexameric fraction was 1.65 nM in 300 µL standard buffer.

## Discussion

Our results demonstrate that nanodisc CNE prevents membrane protein aggregation while maintaining the native protein environment and oligomeric conformation throughout. While all three native gel electrophoresis methods—BNE, high-resolution, detergent-based CNE, and simple, nanodisc-based CNE—allow purifying native protein oligomers, only detergent and nanodisc CNE offer the possibility of confirming the oligomeric state by FCS, as fluorescence quenching does not occur. Importantly, only nanodisc CNE avoids the addition of substances that interact and, possibly, interfere with the protein of interest, that is, only nanodisc CNE avoid the risk of compromising protein conformation and activity. Moreover, nanodisc CNE avoids protein aggregation, one of the most challenging problems in membrane-protein analysis by native gels. Since nanodiscs are used throughout the analysis, the protein of interest is always kept in a native-like lipid-bilayer environment.

Interestingly, this gentle process reproducibly extracts oligomers from the native membrane that have fewer subunits than is generally considered the consensus. We find monomers for the actually tetrameric aqua(glycero)porin GlpF, the tetrameric potassium channel KvAP or the tetrameric sodium channel NavMs and the hexameric *Hp*UreI. An oligomeric distribution upon protein extraction by native nanodiscs has been observed previously ^31^. Using photobleaching, the authors found that the bleaching step distributions for different membrane proteins, among them KcsA, another tetrameric potassium channel, was compatible with a mix of oligomeric states. This illustrates an equilibrium with the complete oligomer. In 2016, this was demonstrated by Chadda et al. measuring the equilibrium free energy of ClC-ec1, a ClC Cl^-^/ H^+^ antiporter in lipid bilayers by diluting the protein into large membranes and quantifying the change in the monomer vs. dimer population utilizing a single-molecule photobleaching analysis ^32^. Similarly, super-resolution localization microscopy revealed that the peptide antibiotic transporter, SbmA, exists in a monomer-dimer equilibrium in *E. coli* ^33^. Furthermore, again using fluorescence microscopy, phosphatidylinositol-4,5-biphosphate (PIP_2_) binding was shown to shift the dynamic equilibrium of the human serotonin transporter, *h*SERT, in the plasma membrane towards a stable dimeric population ^34^. A process defining the protein quaternary structure independent of the protein density at the cell surface. For aquaporins, to the best of our knowledge, reports of monomeric versions are limited to mutated protein variants in artificial membrane systems ^29, 35-37^. There is no evidence for monomeric *Hp*UreI so far. Compared to GlpF ^38^ and *Hp*UreI ^39^, where the functional pores reside within the protomers, monomeric KvAP and NavMs are functionally irrelevant since the pores are formed within the tetramers ^40, 41^. Generally, different oligomeric states of membrane proteins in the plasma membrane can be part of a dynamic equilibrium of the natural folding process of the quaternary protein structure in prokaryotes or a regulatory strategy to tune protein stability, function, and selectivity.

**Table 1.** Summary of molecular brightness, residence times and diffusion coefficients of oligomeric GlpF, NavMs, KvAP and HpUreI, before and after native PAGE.

| Protein | Native PAGE | Molecular brightness [kHz] | Residence time [μs] | Diffusion coefficient [μm<sup>2</sup>/s] |
| --- | --- | --- | --- | --- |

**Table 1. Summary of molecular brightness, residence times and diffusion coefficients of oligomeric GlpF, NavMs, KvAP and HpUrel, before and after native PAGE.**
|  |  | *Before*<br>*native*<br>*PAGE* | *After native*<br>*PAGE* | *Before*<br>*native*<br>*PAGE* | *After native*<br>*PAGE* | *Before*<br>*native*<br>*PAGE* | *After native*<br>*PAGE* |
| --- | --- | --- | --- | --- | --- | --- | --- |
| GlpF | BNE | 33 ±2 | 7.6 ±0.9 | 817 ±85 | 854 ±240 | 32.6 ±3.8 | 33.3 ±8.1 |
| NavMs | hrCNE | 23.4 ±0.4 | 21.5 ±0.9 | 726 ±37 | 740 ±62 | 35.3 ±2.1 | 34.3 ±2.9 |
| KvAP |  | 44 ±0.5 | 48 ±1.7 | 880 ±64.1 | 770 ±107 | 31 ±3.2 | 36.3 ±5.1 |
|  | CNE | 43 ±1.8 | 38 ±1.5 | 747 ±52.6 | 760 ±20 | 36.8 ±2.5 | 35 ±2.6 |
| HpUrel |  | 36 ±1.7 | 37 ±0.6 | 811 ±58.7 | 830 ±84.2 | 34 ±2.4 | 33 ±3.5 |

**Table 2.**
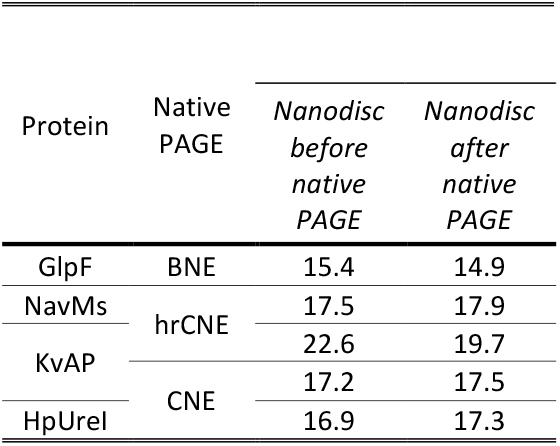
Summary of particle radii of oligomeric GlpF, NavMs, KvAP and HpUreI, before and after native PAGE.

## Conclusion

The combination of gentle membrane-protein extraction into polymer-encapsulated nanodiscs with single-molecule detection methods, such as FCS, advances native PAGE techniques to a new level. It effectively addresses the previously unmet need for substitutes of commercially available membrane-protein weight markers and provides a robust approach to avoid membrane-protein aggregation in the absence of detergents. When combined with fluorescence-labeling of membrane proteins directly in nanodiscs, CNE emerges not only as a reliable quality control method but also as a highly efficient strategy for the purification and analysis of membrane proteins within a native-like lipid-bilayer environment.

## Materials and methods

### Materials

Unless otherwise stated, chemicals were purchased from Sigma–Aldrich or Thermo Fisher Scientific. Glyco-DIBMA polymer (25 g) was purchased from GLYCON Biochemicals, Germany. PureCube 100 Ni-NTA Agarose beads were purchased from Cube Biotech, Germany. ÄKTA Pure chromatography system was purchased from GE Healthcare, U.K. Alexa Fluor 647 maleimide dye was purchased from Jena Biosciences, Germany. Gradient polyacrylamide gels for native PAGE were prepared in-house following protocols. Gel tanks, power supply, and imaging system for native PAGE were purchased from BioRad, UK. MicroTime 200 laser scanning confocal microscope with FLIMbee galvo scanner was purchased from PicoQuant, Germany.

### Protein overexpression and purification

First, an overnight culture supplemented with ampicillin (in 1:1000 dilution) was prepared from 50 mL Lysogeny broth (LB) medium and 1 mL glycerol stock of the protein of interest. The cells were allowed to grow overnight at 37°C, shaking at 180 rpm. The next day, the overnight culture was diluted with 1 L LB medium, supplemented with ampicillin (in 1:1000 dilution). The cells had been shaken at 180 rpm, 37°C until the optical density of the culture reached 0.8; at this point, the cells were inoculated with 1 mL isopropyl-β-D-thiogalactopyranoside (IPTG) to overexpress the protein of interest overnight at 25°C, shaking at 140 rpm. The next day, the cells were collected by centrifugation at 12,000 g, for 10 min. The pellets were resuspended in 40 mL standard buffer (150 mM NaCl or KCl, 50 mM Tris-HCl, pH 8) supplemented with 1 tablet of cOmplete Mini EDTA-free protease inhibitor, 2.5 mM MgSO_4_, and 5 mg/mL DNase. Cells were lysed by French pressure cell press, followed by a 20-min centrifugation at 50,000 g to pellet unlysed cells. Disrupted cells were ultracentrifuged at 100,000 *g*, for 1 h to separate membrane fraction from cytosolic fraction. The membrane fraction was resuspended in the concentration of 50–50 mg/mL and mixed with the same concentration of Glyco-DIBMA (dissolved in the same standard buffer as the C43 cell pellets), in a ratio of 2:1 to 1:2, supplemented with 4 mM MgCl_2_. The solubilization of the membrane fraction happened overnight at 18–20°C. The next day, the sample was ultracentrifuged again (50 000 *g*, 30 min). The His-tagged protein of interest was further purified by Ni^2+^ affinity chromatography and eventually by size-exclusion chromatography.

### Blue and clear native gel electrophoresis

The Ponceau S/glycerol stock solution (0.1% Ponceau S, 50% (*w*/*v*) glycerol was diluted with either the total protein elution or the SEC-separated oligomeric protein material in 1:10 ratio or CNE was performed at 4°C; the power supply was set to 100 V for the first 15–20 min, letting the samples diffuse from the stacking gel into the running gel, then increased the input voltage limited to 160 V. Electrophoresis was continued until the red Ponceau front was 1–2 cm above the bottom of the gel. The gel was fluorescently imaged with Bio-Rad imaging system, using a multichannel excitation program. Fluorescent protein bands were cut out from the gel with a scalpel and resuspended in 200– 300 µL standard buffer overnight at 18–20°C to extract membrane proteins.

*FCS* was used to measure molecular brightness and diffusion coefficient after size-exclusion chromatography (black autocorrelation curves) and after CNE (red and blue autocorrelation curves). We used laser scanning confocal fluorescence microscope to measure the diffusion of Alexa Fluor 647 maleimide labeled protein through the diffraction-limited observation volume. This gave rise to the temporal fluorescence intensity fluctuations that were detected by avalanche photodiodes after having passed through a band pass emission filter ^22, 42^. The measured residence time of the particles in the focal volume, that is, the time it took the particles need to diffuse through this volume, was used to first estimate the diffusion coefficient of the diffusing particle and subsequently the diameter of the nanodisc (see Supplement). The confocal volume was calibrated using Alexa Fluor 647 maleimide dye alone.

## Acknowledgments

This research was funded by the Austrian Science Fund (FWF), grant numbers TAI181, P34826 (both to PP), and P 35541 (to AH).

