## Supplemental Information for "Clear Native Gel Electrophoresis for the Purification of Fluorescently Labeled Membrane Proteins in Native Nanodiscs"

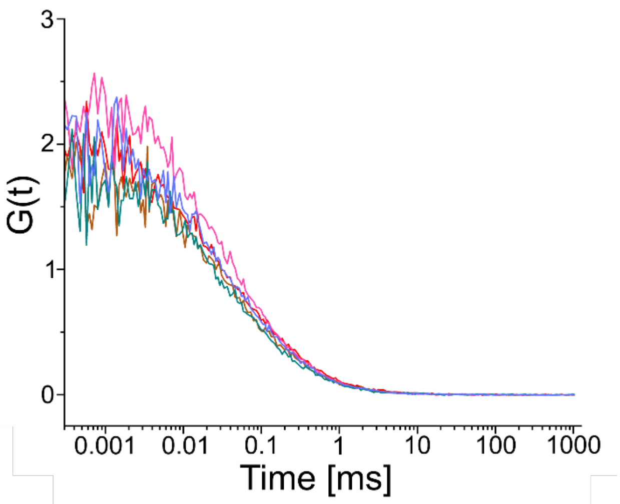


***Figure S1***. 1 nM Alexa Fluor 647 maleimide in solution, measured for molecular brightness and confocal radius and volume estimation before each individual protein measurements: before GlpF/blue native PAGE (red), NavMs/high-resolution CNE (magenta), KvAP/high-resolution CNE (brown), KvAP/detergent-free CNE (cyan) and HpUreI/detergent-free CNE (purple). A 3D extended triplet fit was used to fit the autocorrelation curves.

| | Molecular brightness [kHz] | Residence time [µs] |
|---|---|---|
| GlpF/BNE | 7.5 | 120 |
| NavMs/hrCNE | 6.8 | 85 |
| KvAP/hrCNE | 5.9 | 80 |
| KvAP/CNE | 6 | 89 |
| HpUreI/CNE | 5.8 | 99 |

***Table S1.*** Estimated molecular brightness and residence time of 1 nM Alexa Fluor 647 maleimide dye before the individual protein measurements.

The confocal radius is estimated from the measured residence time and the literature value of the diffusion coefficient of the Alexa Fluor 647 dye (Equation S1):

$$r_0^2 = D * 4 * \tau \qquad \text{(S1)}$$

where $r_0$ is the confocal radius, D is the diffusion coefficient and τ is the residence time.

The effective volume is then estimated [1] (Equation S2):

$$V_{eff} = \pi^{3/2} * r_0^3 * \kappa \qquad \text{(S2)}$$

where $V_{eff}$ is the effective volume and κ is the structural parameter (~8 for 60x objective, 633 laser), which describes the shape of the effective volume.

The concentration is estimated by the following equation (Equation S3):

$$C = \frac{N}{V_{eff} * NA} \qquad \text{(S3)}$$

where C is the concentration, $V_{eff}$ is the effective volume, NA is the Avogadro-constant and N is the number of particles in the effective volume. N is the amplitude of the autocorrelation curve and inversely proportional to G(0).

The average number of lipids per disc is estimated by the subtraction of membrane protein diameter (estimated in Pymol) from the diameter of the nanodisc. This area of lipid is then divided by the area of one lipid head group, which estimated to be roughly 0.7 $nm^2$ in case of prokaryotic membrane lipid mixture.

| Protein | $A_{oligomer}$ [$nm^2$] | $A_{nanodisc}$ [$nm^2$] | $N_{lipid/nanodisc}$ |
|---|---|---|---|
| GlpF | 49 | 174 | 357 |
| NavMs | 28.3 | 222 | 635 |
| KvAP (hrCNE) | 86.6 | 306 | 628 |
| KvAP (CNE) | 86.6 | 240 | 440 |
| HpUreI | 67.9 | 223 | 444 |

***Table S2***. Estimated areas of oligomers and the purified nanodiscs and the estimation of the average number of lipid per nanodisc.


1 Ries, J. & Schwille, P. Studying slow membrane dynamics with continuous wave scanning fluorescence correlation spectroscopy. *Biophysical journal* **91**, 1915-1924, doi:10.1529/biophysj.106.082297 (2006).
